# Fast and interpretable alternative splicing and differential gene-level expression analysis using transcriptome segmentation with Yanagi

**DOI:** 10.1101/364281

**Authors:** Mohamed K Gunady, Stephen M Mount, Héctor Corrada Bravo

## Abstract

**Introduction:** Analysis of differential alternative splicing from RNA-seq data is complicated by the fact that many RNA-seq reads map to multiple transcripts, besides, the annotated transcripts are often a small subset of the possible transcripts of a gene. Here we describe Yanagi, a tool for segmenting transcriptome to create a library of maximal L-disjoint segments from a complete transcriptome annotation. That segment library preserves all transcriptome substrings of length L and transcripts structural relationships while eliminating unnecessary sequence duplications.

**Contributions:** In this paper, we formalize the concept of transcriptome segmentation and propose an efficient algorithm for generating segment libraries based on a length parameter dependent on specific RNA-Seq library construction. The resulting segment sequences can be used with pseudo-alignment tools to quantify expression at the segment level. We characterize the segment libraries for the reference transcriptomes of Drosophila melanogaster and Homo sapiens and provide gene-level visualization of the segments for better interpretability. Then we demonstrate the use of segments-level quantification into gene expression and alternative splicing analysis. The notion of transcript segmentation as introduced here and implemented in Yanagi opens the door for the application of lightweight, ultra-fast pseudo-alignment algorithms in a wide variety of RNA-seq analyses.

**Conclusion:** Using segment library rather than the standard transcriptome succeeds in significantly reducing ambigious alignments where reads are multimapped to several sequences in the reference. That allowed avoiding the quantification step required by standard kmer-based pipelines for gene expression analysis. Moreover, using segment counts as statistics for alternative splicing analysis enables achieving comparable performance to counting-based approaches (e.g. rMATS) while rather using fast and lighthweight pseudo alignment.

## 1 Introduction

Messenger RNA transcript abundance estimation from RNA-Seq data is a crucial task in high-throughput studies that seek to describe the effect of genetic or environmental changes on gene expression. Transcript-level analysis and abundance estimation can play a central role in performing fine-grained analysis studying local splicing events and coarse-grained analysis studying changes in gene expressions.

Over the years, various approaches have addressed the joint problems of (gene level) transcript expression quantification and differential alternative RNA processing. Much effort in the area has been dedicated to the problem of efficient alignment, or pseudo-alignment, of reads to a genome or a transcriptome, since this is typically a bottleneck in the analytical processes that start with RNA-Seq reads and yield gene-level expression or differentially expressed transcripts. Among these approaches are alignment techniques such as bowtie [1], Tophat [2, 3], and Cufflinks [4], and newer techniques such as sailfish [5], RapMap [6], Kallisto [7] and Salmon [8], which provide efficient strategies through k-mer counting that are much faster, but maintain comparable, or superior, accuracy.

These methods simplified the expected outcome of the alignment step to find only sufficient read-alignment information required by the quantification step. Given a transcriptome reference, an index of kmers is created and used to find a mapping between reads and the list of compatible transcripts based on each approach’s definition of compatibility. The next step, quantification, would be to resolve the ambiguity in reads that were mapped to multiple transcripts. Multi-mapping reads are common even assuming error free reads, due to shared regions produced by alternative splicing. The ambiguity in mapping reads is resolved using probabilistic models, such as the EM algorithm, to produce the abundance estimate of each transcript [9]. It is at this step that transcript-level abundance estimation still face substantial challenges that inherently affect the underlying analysis.

The presence of sequence repeats and paralogous genes in many organisms creates ambiguity in the placement of reads. More importantly, the fact that alternatively spliced isoforms share substantial portions of their coding regions, greatly increases the proportion of reads coming from these shared regions and consequently reads being multi-mapped becomes more frequent when aligning to annotated transcripts (Figure 1 A-B). In fact, local splicing variations can be joined combinatorially to create a very large number of possible transcripts from many genes. An extreme case is the Drosophila gene Dscam, which can produce over 38,000 transcripts by joining less than 50 exons [10]. More generally, long-read sequencing indicates that although there are correlations between distant splicing choices [11], a large number of possible combinations is typical. Thus, standard annotations, which enumerate only a minimal subset of transcripts from a gene (e.g. [12]) are inadequate descriptions. Furthermore, short read sequencing, which is likely to remain the norm for some time, does not provide information for long-range correlations between splicing events.

**Figure 1.**
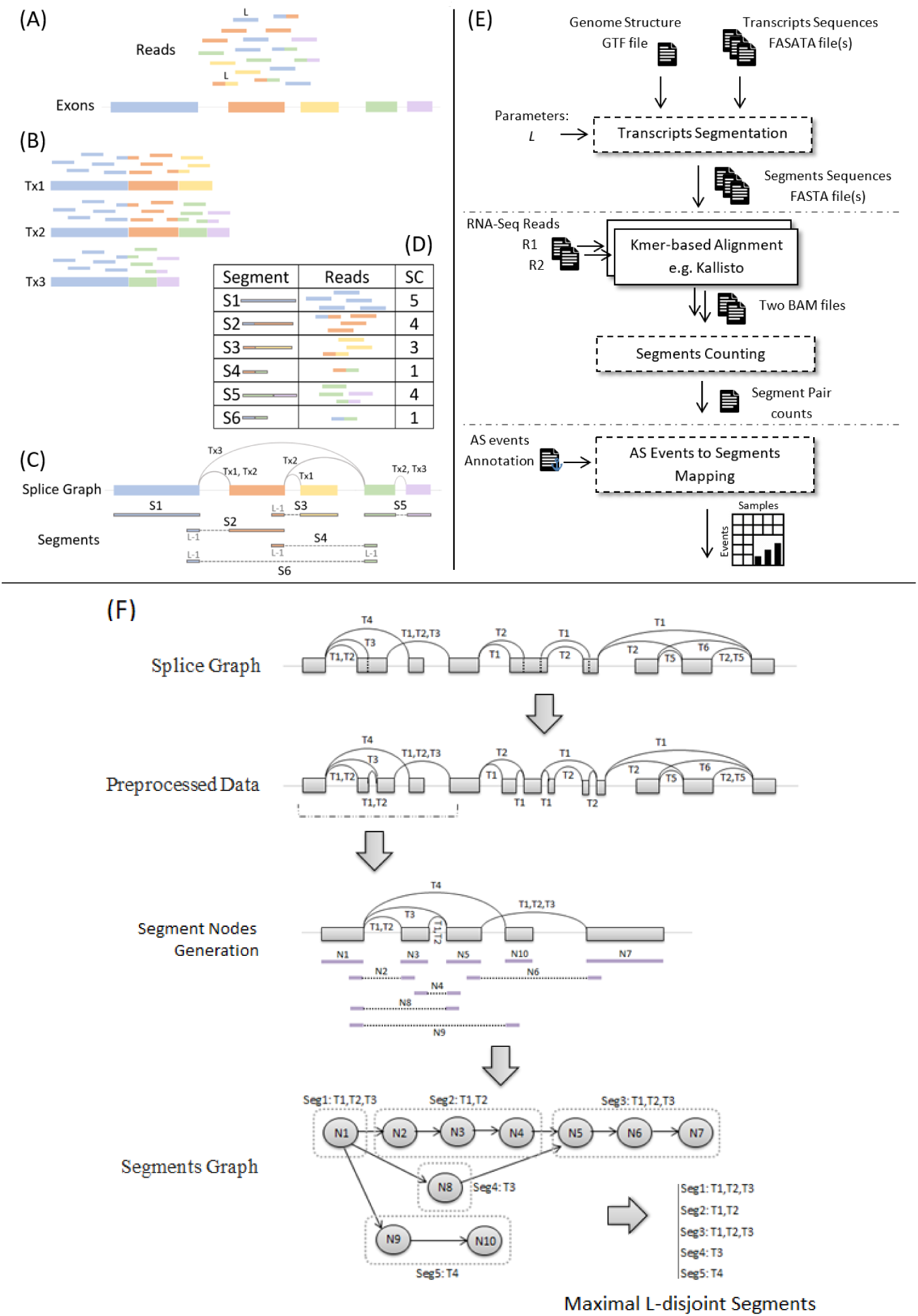
An overview of transcriptome segmentation and Yanagi-based workflow. The leftside shows a typical RNAseq example with Yanagi’s output. (A) Shows the example set of exons and its corresponding sequenced reads. (B) shows the result of alignment over the annotated three isoforms spliced from the exons. (C) shows the splice graph representation of the three isoforms along with the generated segments from yanagi. (D) shows the alignment outcome when using the segments, and its segment counts (SCs). (E) Yanagi-based workflow: segments are used to align a paired-end sample then use the segments counts for downstream alternative splicing analysis. Dotted blocks are components of Yanagi. (F) Yanagi’s three steps for generating segments starting from the splice graph for an example of a complex splicing event. Assuming no short exons for simplicity. Step two and three are cropped to include only the beginning portion of the graph for brevity.

In this paper, we propose a novel strategy that aims at constructing a set of transcriptome segments that can be used in the read-alignment-quantification steps instead of the whole transcriptome without loss of information. Such a set of segments (a segment library) can fully describe individual events (primarily local splicing variation, but also editing sites or sequence variants) independently, leaving the estimation of transcript abundances as a separate problem. Here we introduce and formalize the idea of transcriptome segmentation, propose and analyze an algorithm for transcriptome segmentation, through a tool called Yanagi. To show how the segments library can be used in downstream analysis, we show results of using yanagi for gene-level and alternative splicing differential analysis.

## 2 Transcriptome Segmentation

Figure 1 shows a typical situtation in RNASeq data analysis and provides an overview of the transcript segmentation strategy. In particular, it summarizes how reads that would be multi-mapped when aligning to a transcript library would be aligned to segments. In the latter case, all reads are aligned to a single target sequence and read counts are obtained per segment without the need of probabilistic quantification methods to resolve ambiguity. The next few subsections present a few more specifics of the Yanagi [13] method for transcriptome segmentation.

### 2.1 Segments Properties

Yanagi’s objective is to generate a minimal set of disjoint sequences (where disjointness is parameterized by the experimental sequencing read length) while maintaining transcriptome sequence completeness.

The following definitions are for a given transcriptome T, and parameter L.

#### Definition 1

*A Segment*

*A segment seg defined by the tuple 〈exs, loc, w〉 is a genomic region of width w beginning at genomic location loc and spanning the sequence of consecutive exonic regions exs ∈ Exs_T_ (either exons or retained introns). Exonic regions are considered consecutive if they are consecutively spliced into at least one possible isoform in T. And for all segments in a segments library S_T, L_, its width w is at least L bases*.

#### Definition 2

*Segments Sequences Completeness*

*The set of segments S_T, L_ is Complete if and only if*

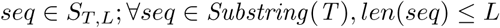

and

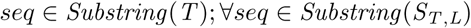

#### Definition 3

*L-disjoint Segments*

*Each segment in the set S_T, L_ is L-disjoint if and only if*

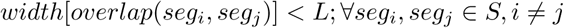

The L-disjointness property restricts any pair of *L-disjoint* segments to have an overlap region shorter than parameter *L*, which typically equals to the sequencing read length. In other words no read of length at least *L* can be mapped to both segments of an *L-disjoint* segment pair, assuming error-free reads.

Another property of the generated segments is to be maximal. For *seg*: 〈*exs*, *loc*, *w*〉, denote *Txs*(*seg*) as the set intersection of annotated transcripts splicing exons *exs*. We can define a subsumption relationship between segments as segi ≻ *seg*_2_ if and only if *exs*_1_ = *exs*_2_, *loc*_1_ = *loc*_2_, *Txs*(*seg*_1_) = *Txs*(*seg*_2_) and *w*_1_ > *w*_2_. With this relationship we can define the following property of a segment library *S*_*T*, *L*_

#### Definition 4

*Maximal Segments*

*For each segment in the set S_T, L_ to be Maximal*

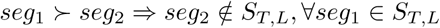

Thus a maximal segment is the longest common sequence of genomic regions starting at *loc*, such that these regions are spliced similarly, i.e. the entire sequence belongs to the same set of transcripts. That is why in figure 1 (C) segment S5 is extended to include two exons and its junction, while segment S2 is interrupted by the different splicings of Tx1 and Tx2.

### 2.2 Segmentation Algorithm

The transcriptome segmentation process can be summarized into three steps: (1) Preprocessing the transcriptome annotation in order to obtain disjoint exonic bins, (2) Constructing a Segments Graph, and finally (3) Generating the final segments. Transactions in Figure 1 (F) represent these three steps.

#### 1. Annotation Preprocessing

Yanagi applies a preprocessing step to eliminate region overlaps present in the transcriptome reference. Parts of an exon (or a retained intron) can be differentially spliced between isoforms either due alternative 3’/5’ splice sites, or transcription start/end sites. For example, splicing the first and second exons between Tx1 and Tx3 in figure 1(F). This step ensures that any splicing event is occurring either at the beginning or the end of an exonic bin, which makes the process of generating maximal L-disjoint segments easier. The preprocessing step is independent from the parameter *L*, so it can be done only once per transcriptome reference.

#### 2. Constructing Segments Graph

Currently Yanagi builds a separate segment graph for each gene, since there are no alternative splicing events between transcripts of different genes. However, future work may use segment graphs that connect different genes sharing regions of identical sequence length L or greater, but we have yet to address this.

##### Definition 5

*Segments Graph*

*A segment graph G_T, L_ is an acyclic directed graph defined by the pair (N, E), where N is a set of nodes representing segments, and E is the set of directed edges between the nodes. An edge e: (n_i_, n_j_) ∈ E is created if the segment corresponding to node n_i_ directly precedes the segment corresponding to node n_j_ in some transcript*.

For each gene, the preprocessed Splice graph is parsed to construct a set of segment nodes (review algorithm details in [13]). These nodes formulate the segments graph of that gene. Each segment node represents an L-disjoint segment, which is not necessarily a maximal segment.

#### 3. Generating Segments

To preserve the maximality property, the segments graph is parsed to aggregated segment nodes into the final maximal segments. In a segment graph, if there is an edge from *node*_*i*_ to *node*_*j*_ while *outdegree*(*node*_*i*_) = *indegree*(*node*_*j*_) = 1, that implies that both nodes belong to the same set of transcripts and can be aggregated into a segment that subsumes both nodes. In other words, aggregating nodes along a path in the segment graph bounded by branching points (nodes with indegree or outdegree greater than 1).

Yanagi reports the segments into a FASTA file. Each sequence represents a maximal L-disjoint segment. Each segment sequence has a header specifying metadata of how each segment was formed, including: gene ID, the set of exonic bins *exs* included in the segment, genome location in the first exonic bin of *exs* where the segment starts, genome location in the last exonic bin of *exs* where the segment ends, and the set of transcripts splicing the segment’s region.

### 2.3 Yanagi-based Workflow

Figure 1 (E) gives an overview of a yanagi-based workflow which consists of three steps. The first step is the transcriptome segmentation, in which the segments library is generated. Given the transcriptome annotation and the genome sequences, and for a specific parameter value *L*, Yanagi generates the segments in FASTA file format. This step of library preparation is done once independently from the samples. The second step is the alignment step. Using any kmer-based aligner e.g. kallisto or RapMap, the aligner uses the segments library for library indexing and alignment. The outcome of this step is read counts per segments (in case of singleend reads) or segment-pair counts (in case of paired-end reads). These segment counts (SCs) are the statistics that yanagi provides to be used in any downstream analysis. The third step depends on the specific target analysis. Later on this work, we describe two use cases where using segment counts shows to be computationally efficient and statistically beneficial.

## 3 Datasets

The experiments are based on the simulation data provided by [14] for both fruit fly and human organisms (dm3 and hg37 assembly versions, respectively). Each dataset consists of samples from two conditions. Each condition has three replicates. The reads for the replicates are simulated from real RNA-seq samples, to get realistic expression values, after incorporating a variance model and the change required between conditions. The simulation is restricted to only protein-coding genes in the primary genome assembly. The difference in transcripts usage across conditions was simulated in 1000 genes randomly selected from genes with at least two transcripts and high enough expression levels. For each of these 1000 genes, the expression levels of the two most abundant transcripts is switched across conditions. Refer to [14] for full details of the preparation procedure of the dataset.

## 4 Analysis of Generated Segments

For practical understanding of the generated segments, we used Yanagi to build segment libraries for the fruit fly and human genomes: Drosophila melanogaster (UCSC dm6) and Homo sapiens (UCSC hg38) genome assemblies and annotations. These organisms show different genome characteristics, *e.g*. the fruit fly genome has longer exons and transcripts than the human genome, while the number of transcripts per gene is much higher for human genome than the fruit fly. A summary of the properties of each genome is found in [14].

### 4.1 Sequence lengths of generated segments

Since *L* is the only parameter required by the segmentation algorithm, we tried different values of *L* to understand the impact of that choice on the generated segments library. Recall that the choice of *L* is based on the expected read length of the sequencing experiment. For this analysis we chose the set *L* = (40,100,1000,10000).

Figure 2 shows the histogram of the lengths of the generated segments compared to the histogram of the transcripts lengths, for each value of *L*, for both fruit fly (left) and human (right) genomes. The figure shows the expected behavior when increasing the value of *L*; using small values of *L* tends to shred the transcriptome more (higher frequencies for small sequence lengths), especially with genomes of complex splicing structure like the human genome. While with high values of *L*, such as *L* = 10, 000, the minimum segment length anticipated tends to be higher than the length of most transcripts, ending up generating segments such that each segment represents a full transcript.

**Figure 2.**
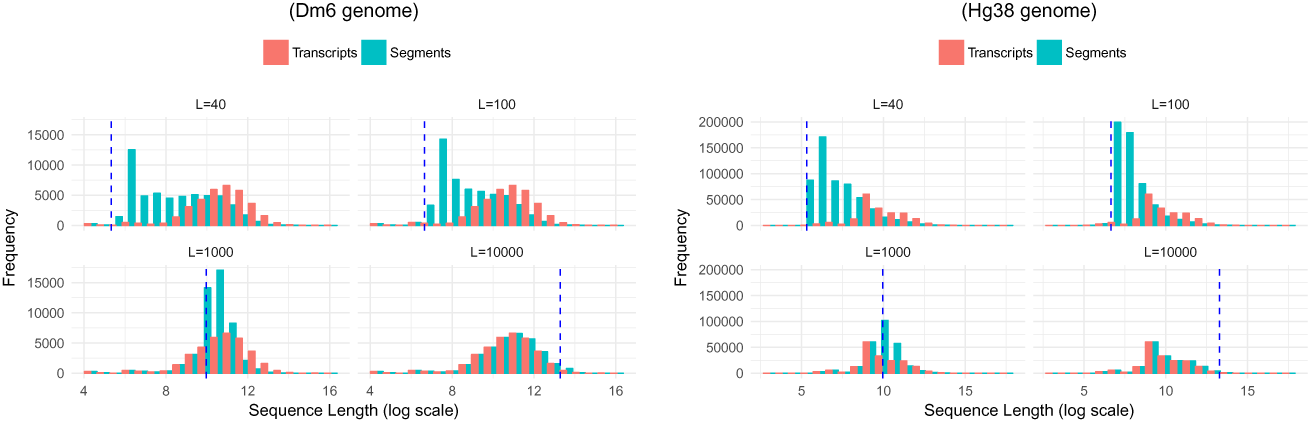
Histogram of transcripts lengths vs. segments lengths for both fruit fly (left) and human (right) genomes, with different values of *L* (40, 108, 1000, 10,000). Dotted vertical line represents the used value of *L* during the transcriptome segmentation.

### 4.2 Number of generated segments per gene

Figure 3 shows how the number of generated segments in a gene is compared to the number of the transcripts in that gene, for each value of *L*, for both fruit fly (left) and human (right) genomes. A similar behavior is observed while increasing the value *L*, as with the segments length distribution. The fitted line included in each scatter plot provides indication of how the number of target sequences grows compared to the original transcriptome. For example when using *L* = 100 (a suitable value with Illumina’s short reads), the number of target sequences per gene, which will be the target of the subsequent pseudo-alignment steps, almost doubles. It is clear from both figures the effect of the third step in the segmentation stage. It is important not to shred the transcriptome so much that the target sequences become very short leading to resulting complications in the pseudo-alignment and quantification steps, and not to increase the number of target sequences leading to increasing the processing complexity of these steps.

**Figure 3.**
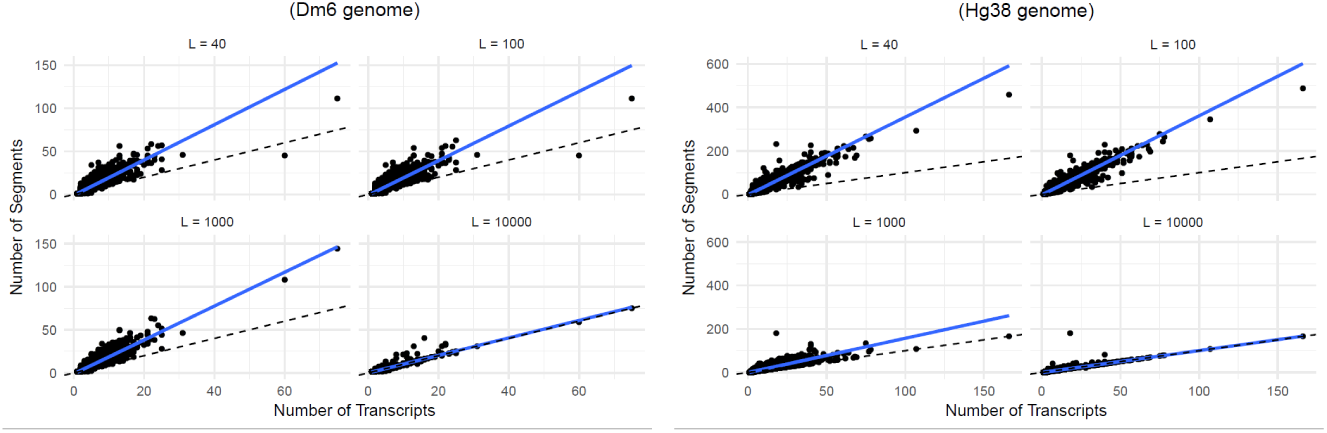
Number of transcripts vs. number of segments, per gene, for both fruit fly (left) and human (right) genomes, with different values of *L* (40, 108, 1000, 10,000). The figure shows how a fitted line (solid blue) compares to the identity line (dotted black).

### 4.3 Library Size of the generated segments

As a summary, Table 1 shows the library size when using segments compared to the reference transcriptome in terms of the total number of sequences, sequence bases, and file sizes. The total number of sequence bases clearly shows the advantage of using segments to reduce repeated sequences appearing in the library that corresponds to genomic regions shared among multiple isoforms. For instance, using *L* = 100 achieves 54% and 35% compression rates in terms of sequence lengths for fruit-fly and human genomes, respectively. The higher the value of *L* is, the more overlap is allowed between segments, hence providing less the compression rate. Moreover, that necessarily hints on the expected behavior of the alignment step in terms of the frequency of multi-mappings.

**Table 1.**
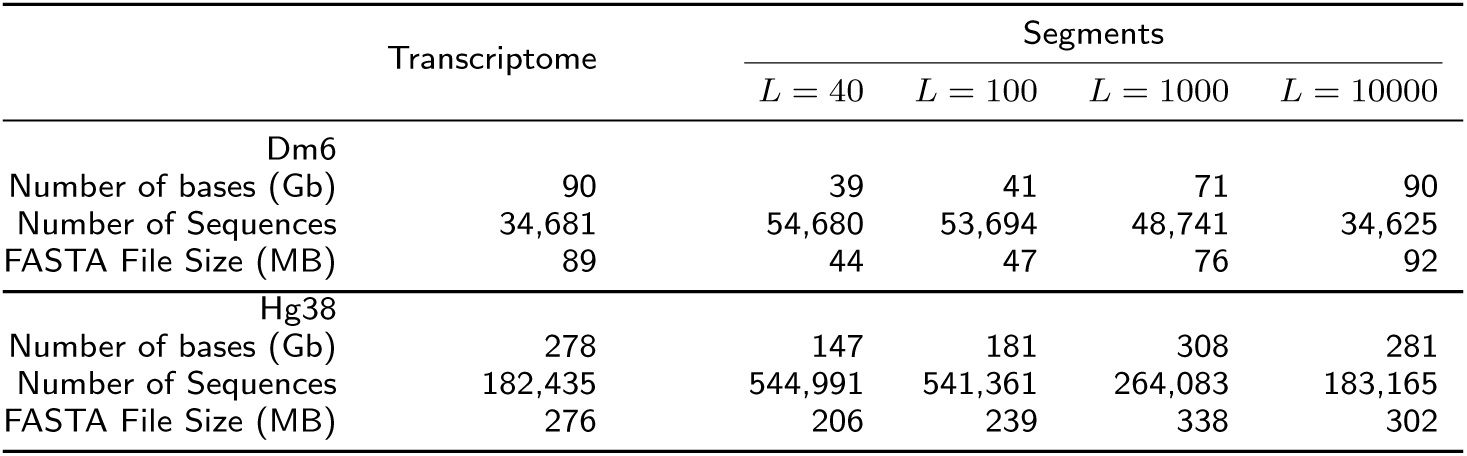
Library size summary

### 4.4 Impact of using segments on Multi-mapped Reads

To study the impact of using the segments library instead of the transcriptome for alignment, we created segments library with different values of *L* and observed the number of multimapped and unmapped reads for each case and how it is compared to when the transcriptome is used. We used RapMap [6] as our kmer-based aligner, to align samples of 40 million simulated reads of length 101 (samples from the dataset discussed in Datasets section) in a single-end mode. The experimented values of *L* were centered around the value of *L* = 101 with more value points close to 101 to test how sensitive the results are towards small changes in the selection of *L*. Figure 4 shows the alignment performance in terms of the number of multimapped reads (red solid line) and unmapped reads (blue solid line), compared to the number of multimapped reads (red dotted line) and unmapped reads (blue dotted line) when aligning using the transcriptome. Using segments highly reduces the number of multimapped reads. The plot shows that too short segments compared to the read length results in a lot of unmapped reads. Consequently, choosing *L* to be close to the read length is the optimal choice to minimize multimappings while maintaining a steady number of mapped reads. It is important to note that the best segments configuration still produces some multimappings. That is a result of the presence of reads sequenced from paralogs and sequence repeats that are not tackled in the current version of yanagi. However, it is clear that using segments can achieve around 10 fold decrease in the number of multimappings.

**Figure 4.**
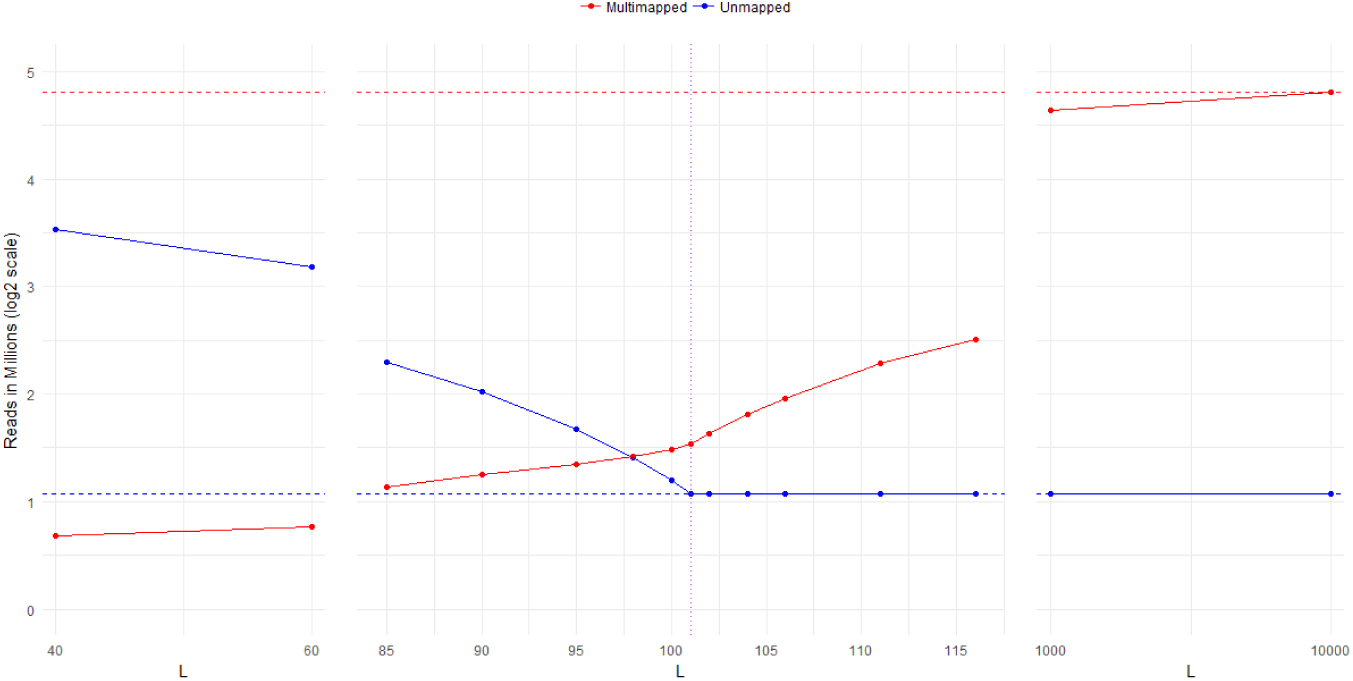
Alignment performance using Segments from hg37, tested for different values of *L*, to align 40 million reads of length 101 (first sample in simulated dataset 3. Performance is shown in terms of the number of multimapped reads (red solid line) and unmapped reads (blue solid line), compared to the number of multimapped reads (red dotted line) and unmapped reads (blue dotted line) when aligning using the transcriptome.

### 4.5 The importance of maximality property

Recalling that the generated segments are maximal segments, as mentioned in definition 2.1. It is the property that segments are extended as much as possible between branching points in the segments graph. The purpose of this property is to maintain stability in the produced segment counts; Since shorter segments will inherently produce lower counts which introduces higher variability that can complicate the downstream analysis. Figure 5 shows the distribution of coefficient of variation (CV) of the produced segment counts from segments with and without maximal property. To examine the effect of the maximal property, we simulated 10 replicates from 1000 random genes (with more than two isoforms) from the hg38 transcriptome using ployester [15]. When segments are created without maximal property, The scatter plot clearly shows that maximal segments have lower CVs to their corresponding short segments for a majority of points (40% of the points has a difference in CVs ¿ 0.05). That corresponds to generating counts with lower means and/or higher variances if the maximal property was dropped.

**Figure 5.**
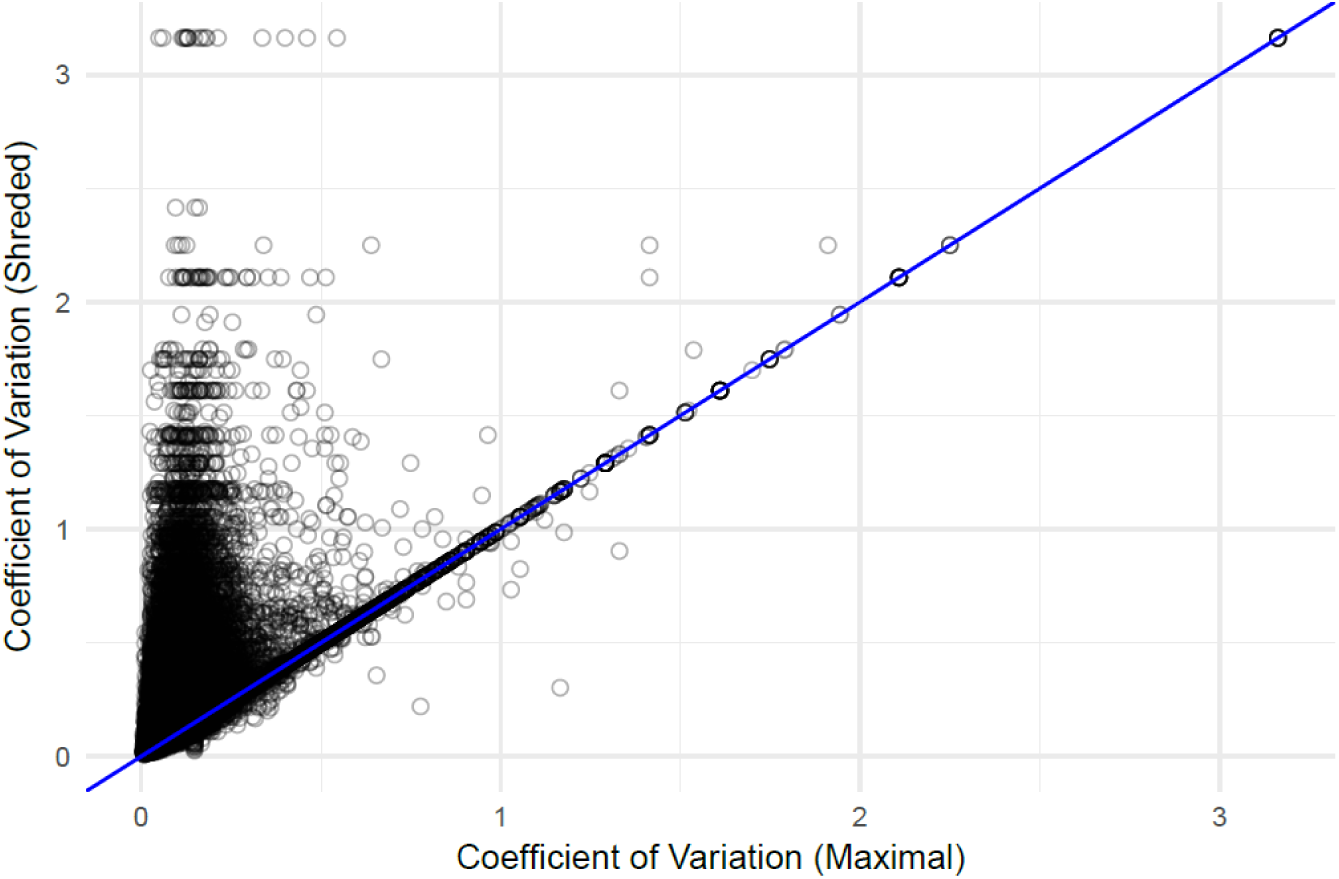
Distribution of coefficient of variation for segment counts produced from maximal segments versus segments without the maximal property enforced. Reads of 10 replicates are simulated from 1000 random genes (with more than two isoforms) from hg38 transcriptome.

## 5 Segment-based Gene Expression Analysis

A typical segment-based approach to do gene expression analysis would start by performing kmer-based alignment over the segments library prepared earlier by Yanagi using high-throughput tools like kallisto, sailfish or RapMap, to derive segment counts (SCs). The segment counts are then used to perform differential gene expression.

The standard RNAseq pipeline for gene expression analysis depends on performing kmer-based alignment over the transcriptome to obtain transcripts abundances, e.g. transcripts per million (TPMs). Then depending on the objective of the differential analysis, an appropriate hypothesis testi is used to detect genes that are differentially expressed. Methods that perform differential gene expression (DGE) prepares gene abundances by summing the underlying transcript abundances. Consequently, DGE methods aims at testing for differences in the overall gene expression. Among these methods are: DESeq2 [16] and edgeR [17]. Such methods fails to detect cases where some transcripts switch usage levels while the total gene abundance is not significantly changing. Note that estimating gene abundances by summing counts from the underlying transcripts can be problematic, as discussed in [18]. RATs [19] on the other hand is among those methods that target to capture such behavior and tests for differential transcript usage (DTU). Regardless of the testing objective, both tests entirely depend on the transcript abundances that were obtained from algorithms like EM during the quantification step to resolve the ambiguity of the multimapped reads, which adherently requires some bias-correction modeling ([8]) adding another layer of complexity to achieve the final goal of gene analysis.

Our segment-based approach aims at breaking the coupling between the quantification, bias modeling, and gene expression analysis, while maintaining the advantage of using ultra-fast pseudo-alignment techniques provided by kmer-based aligners. When Aligning over the L-disjoint segments, the problem of multimapping across target sequences is avoided and as a result the quantification step can be dropped. Then the hypothesis test for differences across conditions are performed on SCs count matrix instead of TPMs.

### 5.1 Kallisto’s TCC-based approach

Yi et al. introduces a comparable approach in [20]. This approach uses an intermediate set defined in kallisto’s index core as equivalence classes (ECs). Specifically, a set of kmers are grouped into an equivalence class (EC) if it belongs to the same set of transcripts during the transcriptome reference indexing step. Then during the alignment step kallisto derives a count statistic for each EC. The statistics are referred to as transcripts compatibility counts (TCCs). In other words, kallisto produces one TCC per EC representing number of fragments that appeared compatible with the corresponding set of transcripts during the pseudo-alignment step. Then the work in [20] uses these TCCs to directly perform gene-level differential analysis by skipping the quantification step using logistic regression. We will refer to that direction as TCC-based approach. To put that approach into perspective with our segment-based approach, we will discuss how the two approaches are compared to each other.

### 5.2 Comparison between segment-based and TCC-based approaches

Both segment-based and TCC-based approaches successfully avoids the quantification step when targeting gene-level analysis. This can be seen as an advantage in efficiency, speed, simplicity, and accuracy, as previously discussed. One difference is that segment-based approach is agnostic to the alignment technique used, while TCC-based approach is a kallisto-specific approach. More importantly, the statistic used in segment-based approach is easily interpretable. Since segments are formed to preserve the genomic location and splicing structure of genes, SCs can be directly mapped and interpreted with respect to the genome coordinates. However, ECs do not have a direct biological meaning in this sense. For instance, all kmers that belong to the same transcript yet originated from different locations over the genome will all fall under the same EC, making TCCs less interpretable. While on the contrary, these kmers will appear in different segments depending on the transcriptome structure. This advantage can be crucial for a biologist who tries to interpret the outcome of the differential analysis. In the next section we show a segment-based gene visualization that allows users to visually examine, for genes determined to be differentially expressed, what transcripts, exons and splicing events contributed to that difference.

Figure 6-bottom shows the number of yanagi’s segments per gene versus the number of kallisto’s equivalence classes per gene. The number of equivalence classes were obtained by building kallisto’s index on hg37 transcriptome, then running the pseudo command of kallisto (kallisto 0.43) on the 6 simulated samplesNote that, in principle there should be more segments than ECs since segments preserve localization, however in practice kallisto reports more ECs than those discovered in the annotation alone in some genes. The extra ECs are formed during pesudo-alignment when reads show evidence of unannotated junctions.

**Figure 6.**
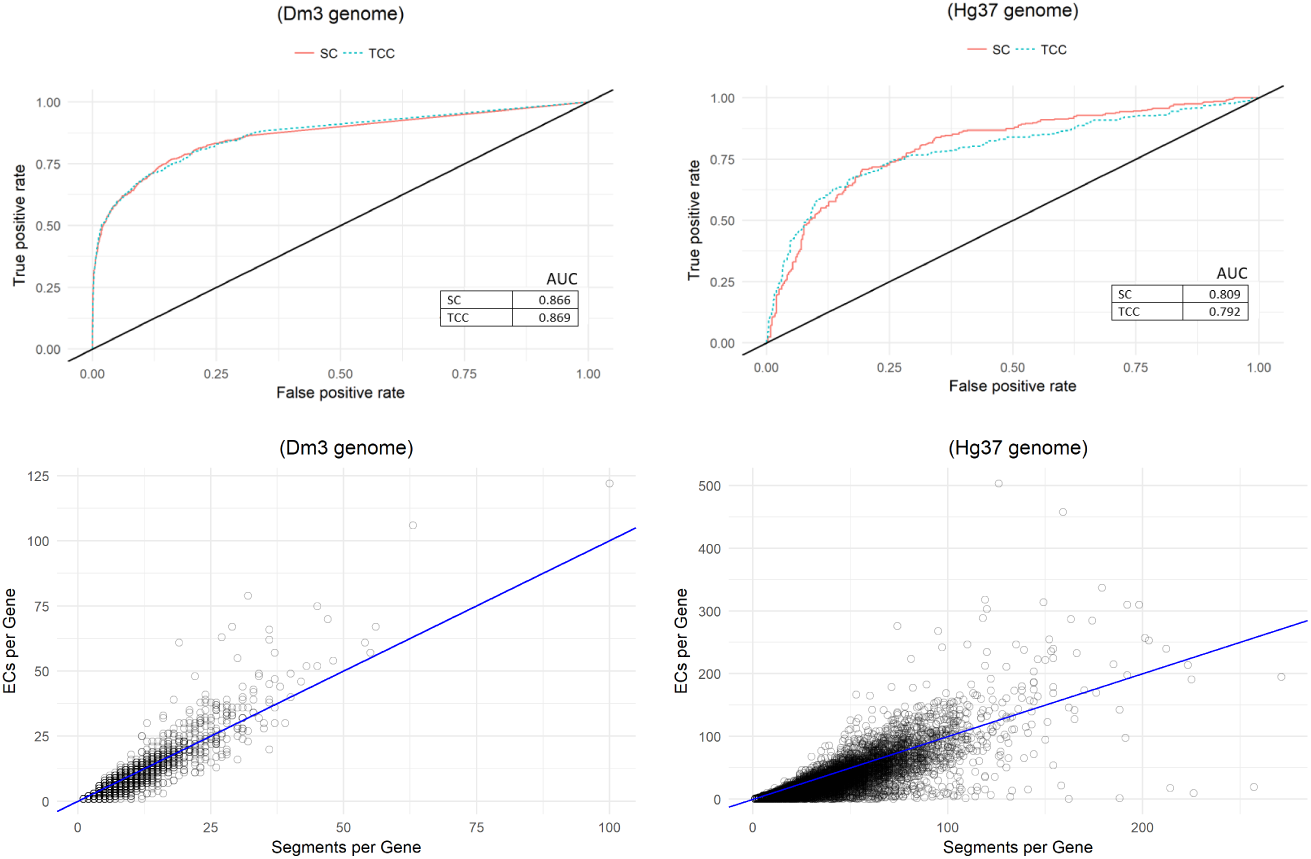
Segment-based gene-level differential expression analysis. Top row, ROC curve for simulation data for DEX-Seq based differential gene-level differential expression test based on segment counts (SC) and Kallisto equivalence class counts (TCC) for *D. melanogaster* and *H. sapiens*. Bottom row, scatter plot of number of segments per gene (x-axis) vs. Kallisto equivalence classes per gene (y-axis) for the same pair of transcriptomes.

### 5.3 DEXSeq-based model for differential analysis

In this work we adopt the DEXSeq [21] method to perform the segment-based gene differential analysis. DEXSeq is a method that performs differential exon usage (DEU). The standard DEXSeq workflow begins by aligning reads to a reference genome, not to the transcriptome, using TopHat2 or STAR [22] to derive exon counts. Then given the exon counts matrix and the transcriptome annotation, DEXSeq tests for DEU after handling coverage biases, technical and biological variations. It fits, per gene, a generalized linear model (GLM) of negative binomial (NB) accounting for effect of the condition factor, and compares it to the null model (without the condition factor) using a chi-square test. Exons that have their null hypotheses rejected are proven to be significanlty different between the experimental conditions, hence DEU is achieved. DEXSeq extends its testing for DEU by controlling the false discovery rate (FDR) at gene level using the Benjamini-Hochberg procedure to find genes with at least one significantly different exon.

Adopting DEXSeq model for the case of segments is done by replacing exons with segments. In other words, the count matrix fed to DEXSeq represent segment counts, instead of exon counts. Once segments are tested for differential usage between conditions, their *p-values* are aggregated to find genes with at least one segment proven to be significantly different.

We tested that model on the simulated data for both human and fruit fly samples, and compared our segment-based approach with the TCC-based approach since they are closely comparable. Since the subject of study is the effectiveness of using either SCs or TCCs as statistic, we fed TCCs reported by kallisto to DEXSeq’s model as well to eliminate any performance bias due the testing model. As anticipated, figure 6-top shows that both approaches provide highly comparable results. Furthermore, using segment counts to test for differentially expressed genes adds to the interpretability of the test outcomes. The next section shows how visualizing segment counts connects the result of the hypotheses test with the underlying biology of the gene.

## 6 Segment-based Gene Visualization

Figure 7 shows Yanagi’s proposed method to visualize segments and the segment counts of a single gene with differentially expressed genes. The plot includes different panels combined, each showing a different aspect of the mechanisms involved in differential expression calls. The main panel of the plot is the segment-exon membership matrix (Panel A). This matrix plot shows the structure of the segments (rows) over the exonic bins (columns) prepared during the annotation preprocessing step. Recall that an exon (or a retained intron) in the genome can be represented with more than one exonic bin in case of within-exon splicing events (Step 1 in section 2.2). Panel B is a transcript-exon membership matrix. It encapsulates the transcriptome annotation with transcripts as rows and the exonic bins as columns. Both membership matrices together allows the user to map segments (through exonic bins) to transcripts.

**Figure 7.**
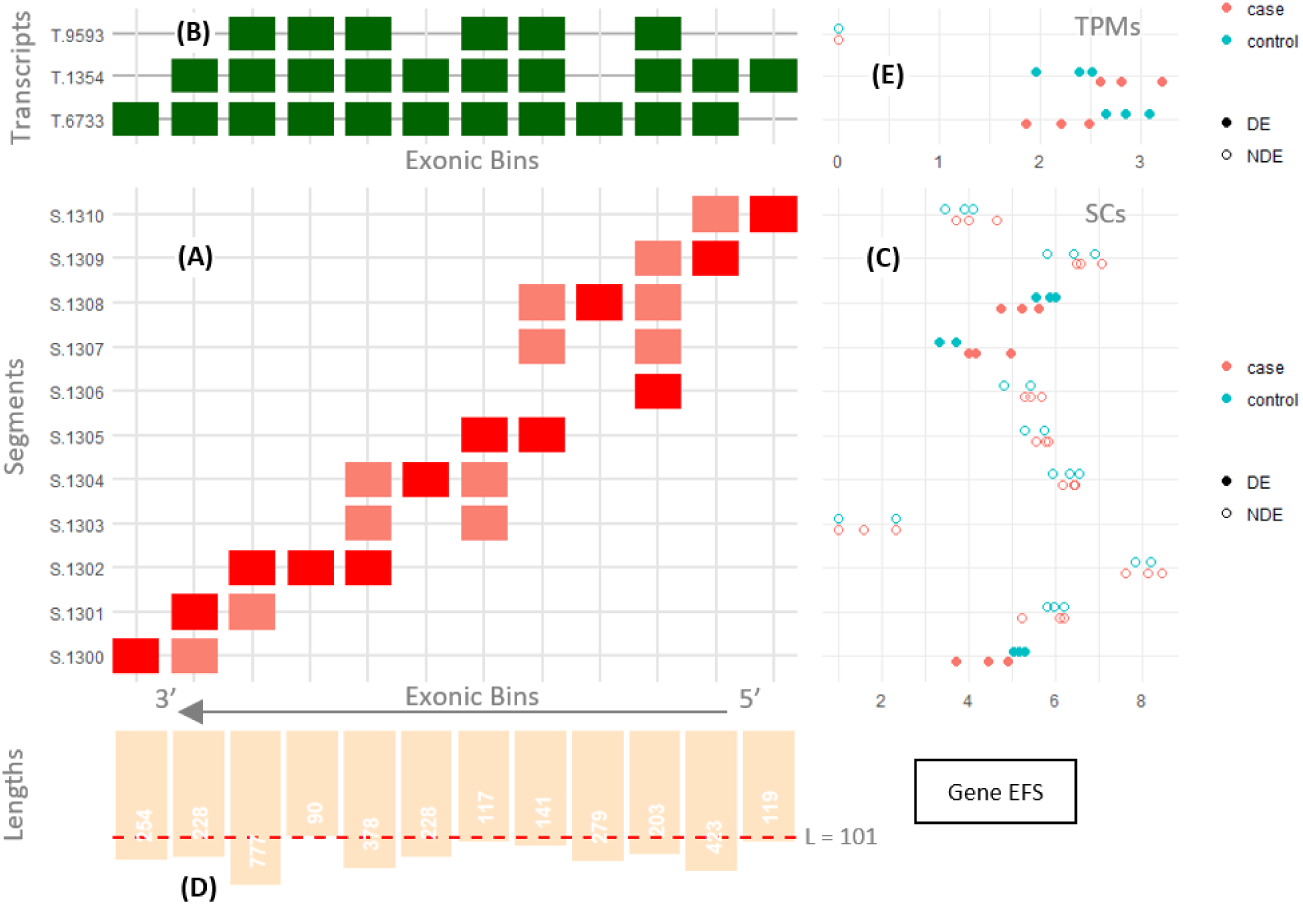
Visualizing segments and segment counts of a single gene with differentially expressed transcripts. It shows human gene EFS (Ensembl ENSG00000100842, genome build Hg37). The gene is on the reverse strand, so the bins axis is reversed and segments are created from right to left. (A) Segment-exonic bin membership matrix, (B) Transcript-exonic bin membership matrix. (C) Segment counts for three control and three case samples, fill used to indicate segments that were significantly differential in the gene. (D) Segment length bar chart, (E) (optional) Estimated TPMs for each transcript.

Panel C shows the segment counts (SCs) for each segment row. Panel D shows the length distribution of the exonic bins. Panel E is optional. It adds the transcript abundances of the samples, if provided. This can be useful to capture cases where coverage biases over the transcriptome is considered, or to capture local switching in abundances that are inconsistent with the overall abundances of the transcripts

The gene in figure 7 is on the reverse strand, that’s why the exonic bins axis is reversed and segments are created from right to left. Consider segment S.0674 for instance. It was formed by spanning the first exonic bin (right-most bin) plus the junction between the first two bins. This junction is present only at transcript T.1354 and hence that segment belongs to only that transcript. In the segment-exon matrix, red-colored cells mean that the segment spans the entire bin, while salmon-colored cells represent partial bin spanning; usually at the start or end of a segment with correspondence to some junction.

Alternative splicing events can be easily visualized from figure 7. For instance, segments S.0672 and S.0671 represent an exon-skipping event where the exon is spliced in T.6733 and skipped in both T.1354 and T.9593.

## 7 Segment-based Alternative Splicing Analysis

Within a gene, the study of how certain genomic regions are alternatively spliced into different isoforms is related to the study of relative transcript abundances. Each local splicing event describes a possible variation of splicing of the described genomic region. For instance, an exon cassette event (exon skipping) describes either including or excluding an exon between the upstream and downstream exons. Consequently, isoforms are formed through a sequential combination of local splicing events. For binary events, the relative abundance of an event is commonly described in terms of percent spliced-in (PSI) [23] which measures the proportion of reads sequenced from one splicing possibility versus the alternative splicing possibilty, while Δ*PSI* describes the difference in PSI across experimental conditions of interest.

Several approaches were introduced to study alternative splicing and its impact in studying multiple diseases. [24] surveyed eight different approaches that are commonly used in the area. These approaches can be roughly categorized into two categories depending on how the event abundance is derived for the analysis. The first category is considered count-based where the approach focuses on local measures spanning specific couunting bins (e.g. exons or junctions) defining the event, like DEXSeq [21], MATS [25] and MAJIQ [26]. Unfortunately, many of these approaches can be expensive in terms of computation and/or storage requirements since it requires mapping reads to the genome, and then processing the huge matrix of counting bins. The second category is isoform-based where the approach uses the relative transcript abundances as basis to derive PSI values. This direction utilizes the transcript abundance (e.g. TPMs) as a summary of the behavior of the underlying local events. Cufflinks [4, 18], DiffSplice [27] and SUPPA [28, 29] are of that category. Unlike Cufflinks and DiffSplice which perform read assembly and discovers novel events, SUPPA succeeds in overcoming the computational and strorage limitations by using transcript abundances that were rapidly prepared by lightweight kmer counting alignment like Kallisto or Salmon.

A main drawback of SUPPA and other transcript-based approaches alike is that it assumes a homogeneous abundance behavior across the transcript making it prone to coverage biases. Previous work showed that RNA-seq data suffers from coverage bias that needs to be modeled into methods that estimate transcript abundances [30, 31]. Sources of bias can vary between fragment length, positional bias due to RNA degradation, and GC content in the fragment sequences. Consider the diagram in figure 8 with a case of two isoforms where isoform1 has higher abundance than isoform2. Both isoforms involve two exon skipping events (E1, E2). The diagram shows the read coverage over different regions of both isoforms with exon E1 in particular has low relative coverage. Considering the real evidence of reads supporting the first skipping event E1, gives a counter conclusion to when considering the overall abundances of the two isoforms involved. More importantly, transcript-based approaches fail to provide different measure of confidence for differential analysis of events E1 and E2 since both events will have the same *PSI* values, whereas there is a significant difference in coverage supporting both events.

**Figure 8.**
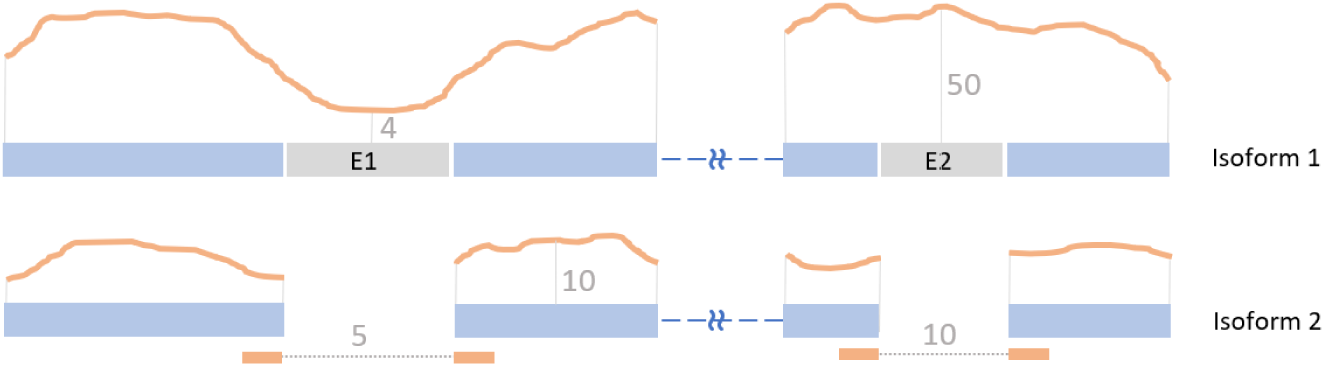
Diagram illustrates the coverage bias problem with AS transcript-based approaches. Given the two given isoforms where isoform1 has higher abundance than isoform2. The diagram shows the read coverage over different regions of both isoforms with exon E1 in particular has low relative coverage. Using the overall transcript abundances gives *PSI* > 0.5 for the first skipping event E1, whereas using the read evidence of the event gives *PSI* < 0.5. Additionally, using transcript abundances gives equal *PSI* values for both events E1 and E2 without any measure of confidence corresponding to their actual evidence.

Our segment-based approach works as a middle ground between count-based and transcript-based approaches. It provides local measures of splicing events while avoiding the computational and storage expenses of count-based approaches by using the rapid lightweight aligners that transcript-based approaches use. Our pipeline begins by running kmer-based lightweight alignment tools like Kallisto over the segments library prepared by Yanagi and obtain the segment counts. Yanagi’s script is then used to map splicing events to their corresponding segments, e.g. each event is mapped into two sets of segments: The first set spans the inclusion splice, and the second for the alternative splice. Current version of Yanagi follows SUPPA’s notation for defining a splice event and can process seven event types: Skipping Exon (SE), Retain Intron (RI), Mutually Exclusive Exons (MX), Alternative 5’ splicec-site (A5), Alternative 3’ splicec-site (A3), Alternative First Exon (AF) and Alternative Last Exon (AL).

### 7.1 Segment-based calculation of PSI

While Yanagi uses the transcriptome annotation to prepare the segments along with the splicing events, it generates mapping between each event and its corresponding segments spanning the event. For each event, Yanagi takes into consideration the transcripts involved and the event genomic coordinates to decide the set of transcriptome segments that correspond to each of the two possibilities of the splicing event. This step becomes complicated in case of overlapping events. The current version of Yanagi selects segments that spans either the event exon or junctions while the segment belong to at least one transcript that undergoes the corresponding splicing.

After alignment, Yanagi provides segment counts or segment-pair counts in case of paired-end reads. For each splicing event, we calculate the PSI value of event *e* in sample *x* as follows:

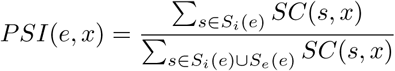

where *S*_*i*_(*e*) and *S*_*e*_(*e*) are inclusion and exclusion segments, respectively, and *SC*(*s*, *x*) is the segment count in the sample. That means segment-based PSI values uses reads spanning both the junctions and the target inclusion exon towards the inclusion count. In fact, read counts will also include reads extended around the event as long as the segment extends on both sides. This extension takes advantage of situations where splicing events are near to include as much discriminative reads into the counts to achieve higher levels of confidence when calculating PSI values.

### 7.2 PSI comparison on simulated data

we compared PSI values obtained from our approach versus counting-based approaches like rMATS and isoform-based approaches like SUPPA2 on splicing events found in hg37 based on the six samples in section 3. Since each tool provides different set of events, We focus our comparison on the intersection set of events between SUPPA and rMATS. That includes events from five types of splicing events. Table 3 summarizes the number of events subject to the study. Two levels of filtering are applied to observe how the different approaches behave in different scenarios. Nonoverlapping events is the smallest subset of events where there is no more splicing other than the two possibilities defining the event, i.e. complex splicing is excluded. While highTPM events is a subset of events in which inclusion and exclusion isoform levels are relatively high (*TPM*_*inc*_ > 1, *TPM*_*ex*_ > 1). This is a typical filtering criteria adopted by isoform-based approaches. This filter excludes events involving isoforms of low levels of expression which inherently suffer from low estimation accuracy.

**Table 2.**
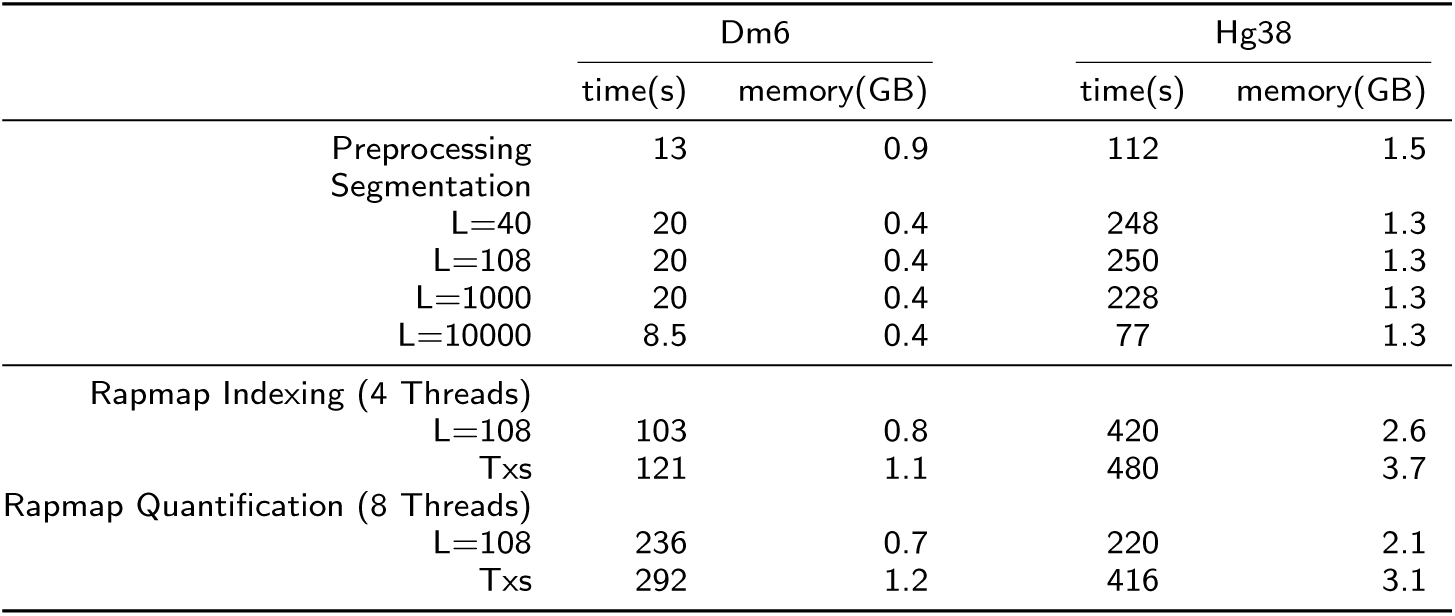
Running time (seconds) and memory usage (gigabytes) by Yanagi to generate segment library for fruit fly (Dm6) and human (Hg38) genomes, for both the preprocessing and segmentation steps. Time for the preprocessing step does not include the time to load the FASTA and GTF files. Most of the memory usage is from loading the input data in both steps. Running on a 6-core 2.1 GHz AMD processor, using single-threaded processes. The lower half shows the time and memory usage for running Rapmap’s quasi-mapping using the segments library and the the full transcriptome, to quantify samples of 40M paired-end reads, each of length 101bp.

**Table 3.** Number of Events in Hg37 common between MATS and SUPPA for the five event types reported by both tools. Two levels of filtering are applied to obtain three subsets. Non-overlapping events are the simplest events where there is no more splicing other than the two possibilities defining the event. While highTPM events are events where inclusion and exclusion isoform levels are relatively high (*TPM*_*inc*_ > 1, *TPM*_*ex*_ > 1).

| Events Subset | SE | MX | A3 | A5 | RI | Total |
| --- | --- | --- | --- | --- | --- | --- |
| Non-overlapping | 4,180 | 68 | 1,435 | 885 | 323 | 6,891 |
| HighTPM Events | 9,756 | 354 | 2,327 | 1,483 | 793 | 14,713 |
| All Events | 13,650 | 1,024 | 3,131 | 2,053 | 1,711 | 21,569 |

Figure 9 shows a scatter plot of PSI values calculated by the three approaches. It is clear that our segment counts (SCs) based approach produces results comparable to rMATS with average Pearson correlation of 0.92 over the full set of events. As expected, PSI values obtained by our approach and rMATS are more correlated to each other than to values derived directly from TPMs, since both our approach and rMATS’s are counts based.

**Figure 9.**
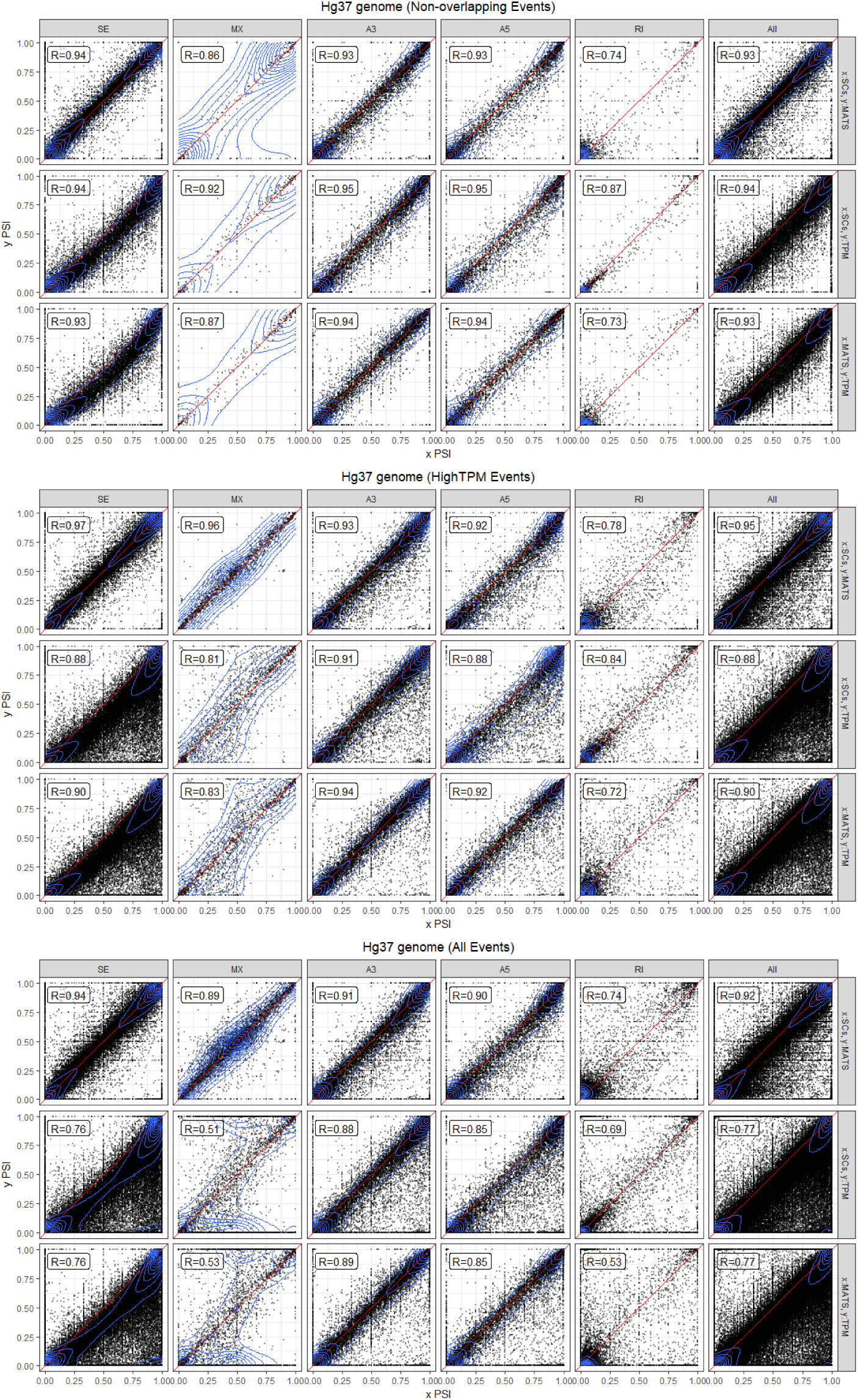
PSI value comparison between segment counts, rMATS (based on spliced alignment to genome) and SUPPA2 (based on estimated TPMs from pseudo-alignemnt and quantification). Columns indicate seven types of alternative splicing events. Scatterplots are stratified by event types (non-overlapping, high TPM, and all events). See Table 3 for number of events of each AS event type shown.

Our results and rMATS are consistently comparable across the three subsets of events. In other words, both approaches give comparable results for cases of events with complex overlapping splicing, While results start to diverge from isoform-based results for overlapping events. On the other hand, results from isoform-based start to be less correlated with the other two approaches when events with low TPMs are included.

Among the five different splicing types exon skipping, alternative 3’ and alternative 5’ events gives the highest correlation between segment counts and rMATS approaches. In our experiments we noticed that rMATS (v4.0.1) does not behave as intended for intron retention events. We noticed that counts including junction reads only and counts including both junction and intron reads (which we use in this study) are the same. In other words, rMATS fails to report reads spanning the intron, which explains the underestimated inclusion counts and PSI values for retained introns.

### 7.3 Differential Alternative Splicing

Since the scope of this paper is to introduce the use of segment counts as a statistic for studying alternative splicing, we want to use the simplest statistical model for differential splicing to exclude any advantage of the model itself. In that matter we used the PSI values of the three approaches (SCs, rMATS, TPM) as discussed in the previous section. Then we used a linear model for differential hypothesis testing (implemented with Limma-voom R Package [32, 33]). However, more advanced models of differential analysis can be used instead. For example, a similar model to SUPPA2 can be developed to test the significance of Δ*PSI* by considering all events genome-wide [29]. Figure 10 shows ROC plots for sensitivity and specificity measures. Using segment counts achieves comparable performance to both rMATS and isoform-based approaches.

**Figure 10.**
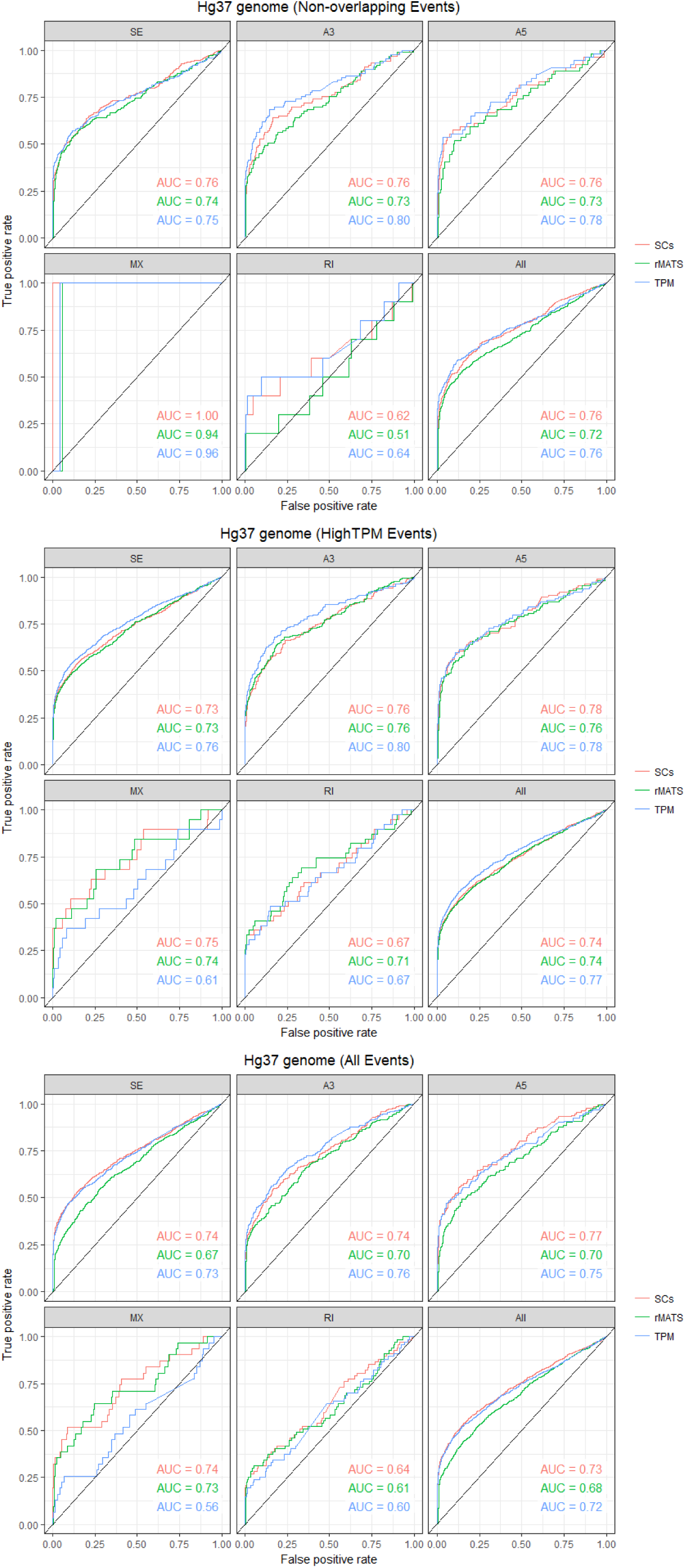
ROC curves for differential alternative splicing comparison between segment-count method, rMATS and SUPPA2 based on simulation data. See Table 3 for number of events of each AS event type shown.

## 8 Discussion

In this paper, we formalized the concept of transcriptome segmentation and propose an efficient algorithm for generating segment libraries based on a length parameter dependent on specific RNA-Seq library construction. The resulting segment sequences were used with pseudo-alignment tools to quantify expression at the segment level. We characterized the segment libraries for the reference transcriptomes of Drosophila melanogaster and Homo sapiens and provided gene-level visualization of the segments for better interpretability. We demonstrated the use of segments-level quantification into gene expression and alternative splicing analysis. The notion of transcript segmentation as introduced here and implemented in Yanagi opens the door for the application of lightweight, ultra-fast pseudo-alignment algorithms in a wide variety of RNA-seq analyses.

Using segment library rather than the standard transcriptome succeeds in sig-nificantly reducing ambigious alignments where reads are multimapped to several sequences in the reference. That allowed avoiding the quantification step required by standard kmer-based pipelines for gene expression analysis. Moreover, using segment counts as statistics for alternative splicing analysis enables achieving comparable performance to counting-based approaches (e.g. rMATS) while rather using fast and lighthweight pseudo alignment that are several folds faster than standard counting pipelines.

## Author’s contributions

HCB, SMM, and MKG conceived and designed the study. HCB and MKG designed all software tools used in the study, MKG implemented all software tools used in the study. MKG acquired all data used and analyzed and interpreted results with HCB. MKG wrote the intial draft of the manuscript, HCB and SMM participated the final drafting of the manuscript.

## Acknowledgements

We want to thank Julien Buchbinder and Steffen Cornwell for preliminary work in transcript segmentation. This work was partially supported by NSF grant ABI 1564785 to SMM and MKG, and NIH grants HG005220 and GM114267 to HCB and MKG.

## Additional Files

Additional file 1 — Sample additional file title

Additional file descriptions text (including details of how to view the file, if it is in a non-standard format or the file extension). This might refer to a multi-page table or a figure.

Additional file 2 — Sample additional file title

Additional file descriptions text.

